# Quantifying 35 transcripts in a single tube: Model-based calibration of the GeXP RT-PCR assay

**DOI:** 10.1101/159723

**Authors:** Pauline Marquardt, Britta Werthmann, Viktoria Rätzel, Markus Haas, Wolfgang Marwan

## Abstract

Quantitative analysis of differential gene expression is of central importance in molecular life sciences. The Gene eXpression Profiling technology (GeXP) relies on multiplex RT-PCR and subsequent capillary electrophoretic separation of the amplification products and allows to quantify the transcripts of up to approximately 35 genes with a single reaction and one dye. Here, we provide a kinetic model of primer binding and PCR product formation as the rational basis for taking and evaluating calibration curves. With the help of a purposeful designed data processing workflow supported by easy-to-use Perl scripts for calibration, data evaluation, and quality control, the calibration procedure and the model predictions were confirmed and the robustness and linearity of transcript quantification demonstrated for differentiating *Physarum polycephalum* plasmodial cells. We conclude that GeXP analysis is a robust, sensitive, and useful method when the transcripts of tens to few hundred genes are to be precisely quantified in a high number of samples.

## Introduction

The quantitative analysis of differential gene expression is of central importance in cell, developmental, and systems biology. Established experimental approaches have specific strengthes and weaknesses and the method of choice depends on the required accuracy for quantitative detection, the available amount of RNA per sample, the number of different transcripts that are to be quantified in one sample, and the available budget [1, 2]. Transcript quantification by the next generation sequencing RNAseq method, for example, allows to quantify the entire transcriptome with an accuracy depending on the number of reads analysed. However, the quantification of one sample is relatively expensive in terms of sequencing costs and requires sophisticated bioinformatic data processing [3, 4]. Real-time quantitative RT-PCR on the other hand is relatively inexpensive and easy to perform but only gives the transcript concentration for upto few genes per sample. A different, appealing, and useful approach is the GeXP multiplex-RT-RCR developed by Beckman-Coulter^1^. This method relies on the simultaneous amplification of the cDNAs of a moderate number of genes in the same sample and the subsequent separation of the fluorescence tagged amplification products by capillary electrophoresis. The number of transcripts that can be quanitfied in a single pot is limited by the number of fragments that can be thoroughly separated from each other. With the CEQ 8800™ Genome Analysis System, an 8-capillary sequencer, at least 35 different transcirpts can be readily quantified using a single fluorescent dye [5].

The method is depicted in Fig. 1. Each transcript of interest is reverse transcribed with a gene-specific RT-primer (reverse primer) and the second strand of the cDNA is synthesized with a gene-specific forward primer. Both, the forward and the reverse primers carry additional sequence stretches at their 5’-ends that are identical for all gene-specific primer pairs used in the multiplex RT-PCR reaction.

**Fig. 1.**
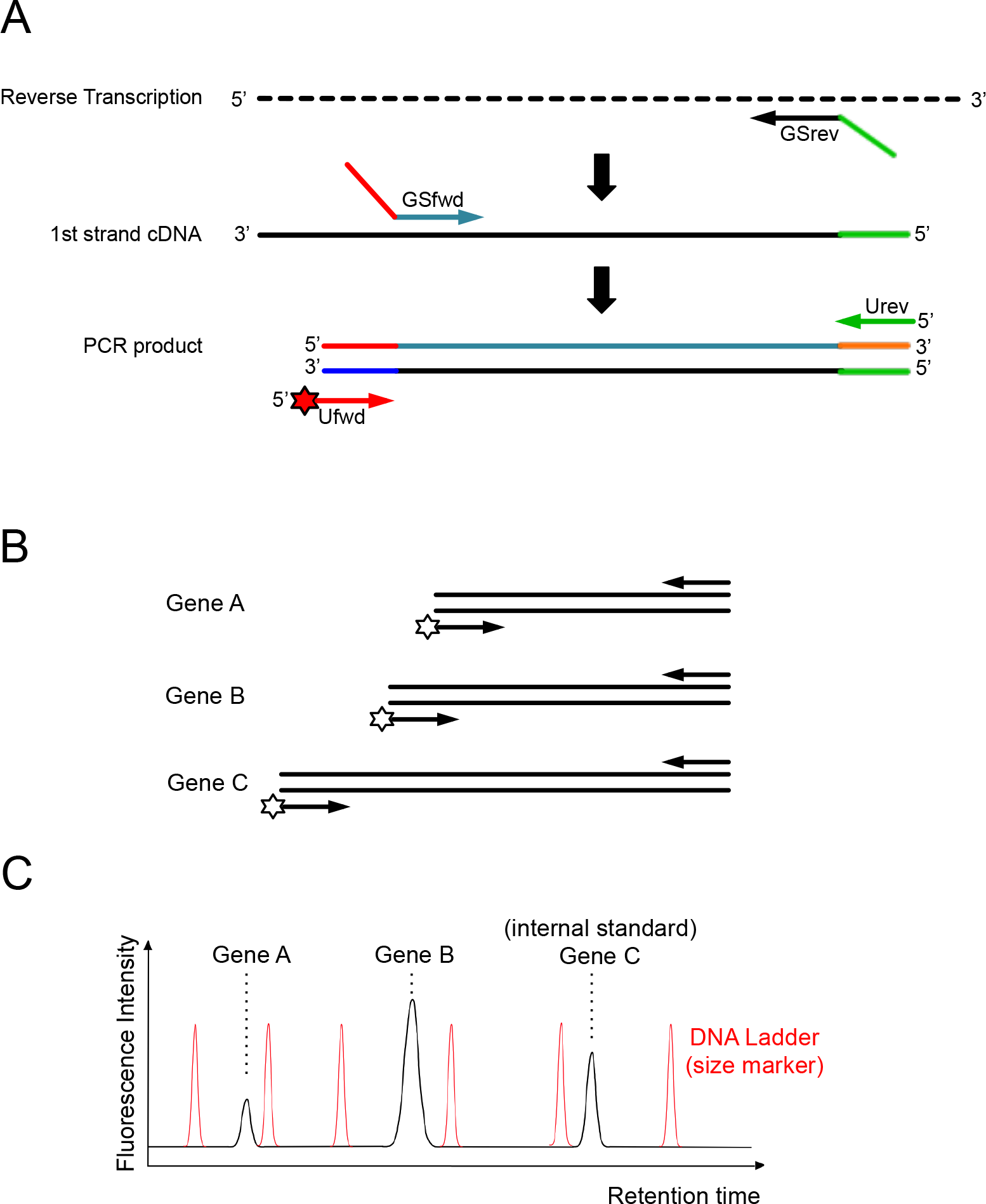
Schematic representation of the GeXP multiplex RT-PCR assay for quantification of relative abundances of transcripts. (A) The reverse transcription of a RNA is primed with a gene-specific reverse primer (GSrev) that carries a universal priming site at its 5’-end to obtain a first strand cDNA fused with the universal priming site. The synthesis of the second strand cDNA is primed with a gene-specific forward primer (GSfwd) that also carries a yet different sequence as universal priming site. Because of the two universal priming sites incorporated, the cDNA can be amplified with universal forward and reverse PCR primers (Ufwd, Urev, respectively) independently of the gene-specific sequence. As the universal forward primer is labelled with a fluorescence tag at its 5’-end, the PCR product can be detected after capillary electrophoretic separation. (B) Gene-specific primers are designed to yield PCR products of different lengths that can be separated by capillary electrophoresis from each other and quantified through the area of individual peaks in the elution profile (C). (C) The gene expression pattern of different samples can be compared through the cDNA amplified from a spike-in RNA that serves as an internal standard. A DNA ladder (size marker) added to the cDNA sample prior to capillary electrophoresis allows to determine the size of the PCR products.

During PCR these sequences serve as specific binding sites for the two universal primers. The purpose of the universal primers is to amplify all cDNAs in the reaction mix independent of their length or sequence with equal efficiency during each cycle of the PCR. With the help of a spike-in RNA, an internal standard which is reverse transcribed together with the RNAs to be analysed, the amount of cDNA fragment amplified from each gene of interest can be normalized relative to the amount of fragment amplified from the spike-in RNA so that the relative transcript abundance for each gene becomes comparable between samples. By producing fragments of different length that are separated electrophoretically (Fig. 1), the relative abundances of the 35 chosen transcripts can be determined provided the system is calibrated appropriately. The normalized peak area for each amplification product which is determined during the electrophoretic separation can be fitted to the total RNA concentration by linear regression of the data points on a double logarithmic plot [6, 7]. Here, we demonstrate that this empirical calibration procedure complies with a kinetic model of cDNA synthesis and amplification. The kinetic model predicts for a set of different transcripts a series of curves of equal slope shifted in parallel to each other. We experimentally confirm the model predictions and the validity of the model-based calibration procedure. Finally, we apply these methods to estimate the relative concentration of different transcripts of differentiation marker genes expressed in *Physarum polycephalum* plasmodial cells that are committed to sporulation [6], see SI Table 1. We also provide a set of Perl scripts as components of an integrated workflow for efficient data processing and evaluation, which is demonstrated in the context of the case study.

## Materials and methods

### Growth and preparation of sporulation-competent plasmodia

Plasmodia of the apogamic strain WT31 [8] which is wild-type with respect to the photocontrol of sporulation were grown for four days at 24 °C in 1.5 L of growth medium [9] in a 5 L fermentor (Minifors, Infors HT, Bottmingen, Switzerland) inoculated with 2 % of a 3.5 days old shaken culture, supplied with 1 L of air per minute, and stirred at 250 rpm with a marine propeller. Microplasmodia were harvested and applied to starvation agar plates (9 cm diameter) as described [8]. Plates were incubated at 22 °C in complete darkness for 6 days if not stated otherwise. During this time period, one multinucleate macroplasmodium develops on each plate and, by starvation, becomes competent for sporulation. The incubation temperature is critical in order to avoid unwanted spontaneous sporulation. For the induction of sporulation, competent plasmodia were exposed to a far-red light pulse (*λ* ≥700 nm, 13 W/m^2^). The far-red light was generated by Concentra Weißlicht lamps (Osram, Munich, Germany) which was passed through an Orange 478 combined to a Blue 627 plexiglass filter (Röhm, Darmstadt, Germany). After light exposure, plasmodia were returned to the dark and incubated at 22 °C until the samples were taken. At five subsequent time points, a portion of each plasmodium was harvested (approximately one sixth of its original biomass for each sample), shock-frozen in liquid nitrogen and stored at −80 °C for RNA isolation. After taking the samples, the remainder of each plasmodium was maintained overnight in the dark to reveal the developmental decision (*i.e.* whether or not sporulation had occurred). Control plasmodia were treated identically except that the light pulse was omitted (dark controls). All manipulations were done under sterile conditions and under dim green safe light as previously described [8].

### Preparation of total RNA

For extraction of total RNA, approximately 15 mg of fresh weight material which had been taken from an individual plasmodium was used. Samples were dissolved in 1.5 ml PeqGOLD TriFastTM solution (Nr 30-2020; PeqLab, Erlangen) by disrupting the cell material in the presence of glass beads ø=0.5 mm using a Precellys 24-Dual homogenizer, (Peqlab, Erlangen, Germany) for 10 seconds at room temperature. The homogenate was transferred to 2 ml PeqGOLD PhaseTrap tubes (PeqLab; Nr 30-0150A-01). After addition of 300 μl CHCl_3_, vigorous shaking and centrifugation of the PhaseTrap tubes, the aqueous phase containing the RNA was transferred to a fresh Eppendorf cap following the manufacturer's instructions for the entire procedure. After addition of 0.9 volumes of a 4:1 mixture of ethanol and 1 M acetic acid, the RNA was precipitated at -20 °C for at least 24 h. The precipitate was spun down for 15 min at 18,000 × g and 4 °C, the pellet washed once with 400 μl ice-cold 3 M sodium acetate and twice with ice-cold 70 % ethanol followed by centrifugation for 10 min at 18,000 × g, 4 °C after each washing step. Finally, the RNA was dissolved in RNAse-free water. For removal of contaminating genomic DNA, 90 μl of each RNA sample were mixed with 10 μl rDNase buffer and 1 μl rDNase (rDNase Set Macherey-Nagel: 740963) and incubated for 12 min at 37 °C. The total RNA was finally purified using the RNA Clean-up Kit (Macherey-Nagel: 740948.250). Approximately 300 ng aliquots checked for integrity of the 18 S and 28 S rRNA by agarose gel electrophoresis. For subsequent steps, samples were standardized to 20 ng/μl of total RNA.

### Multiplex Gene Expression Profiling (GeXP)

Multiplex RT-PCR reactions were performed to simultaneously amplify 35 transcripts and the amplification products were quantified through separation on a Beckman Coulter 8-capillary sequencer (CEQ 8800). The GenomeLab GeXP Start Kit (Beckman Coulter; A85017) along with the ThermoStart Taq Polymerase Kit (Beckman Coulter; A25395) was used according to the GeXP Chemistry Protocol (The Beckman Coulter Capillary Electrophoresis product line including GeXP chemistry is now distributed and supported by AB SCIEX, Landwehrstr. 54, 64293 Darmstadt, Germany (www.SCIEX.com)). cDNA synthesis and polymerase chain reaction were performed in half of the recommended volume leaving the recommended concentrations unchanged. Twenty ng of total RNA was used for each reverse transcription (see below). Controls without template were included to check the kit components for contaminating nucleic acids and controls without reverse transcriptase were use to check for contaminating DNA in the RNA samples.

***Primers used.*** Forward and reverse gene-specific primers for a set of 35 genes were originally desinged by [6] to generate amplified DNA fragments with similar GC contents and melting temperature of different lengths ranging between 114 and 357 bp (SI Table 1). Each primer contained a universal priming sequence at the 5’- end and a gene-specific sequence at the 3’-end. The concentration of each gene-specific reverse primer in the pre-mixed reverse primer plex was experimentally adjusted to give a fluorescence signal for each fragment that was in the linear sensitivity range of the measurement system. The attenuated concentration of each reverse primer depended on its sequence and was adjusted to a value between 0.5 to 0.00025 μM (SI Table 1).

***cDNA synthesis.*** The RT reaction was performed in 96-well plates in a Biometra (T Professional) thermal cycler with 20 ng of total RNA (except for the standard curves), 1 × RT Buffer, 1 μL of reverse primer mix, 1 μL KAN^r^ RNA and 0.5 μL reverse transcriptase (20 U/μl) in a total volume of 10 μL with the following temperature program: 1 min at 48 °C, 5 min at 37 °C, 60 min at 42 °C, 5 min at 95 °C, hold at 4 °C. The plate was briefly centrifuged to re-collect the sample volume at the bottom of the wells.

***Polymerase chain reaction.*** 4.7 μL of the 10 μL RT reaction were used as template for the PCR. The PCR was performed with 0.35 μL Thermo-Start DNA Polymerase, 5 mM MgCl_2_, 1 × PCR Buffer and pre-mixed forward primer plex (20 nM final concentration) in a volume of 10 μL under the conditions: 10 min at 95 °C, followed by 35 cycles of 30 s at 94 °C, 30 s at 55 °C, and 1 min at 68 °C; hold at 4 °C.

***Fragment analysis.*** Amplification products were separated and quantified according to the manufacturer’s instructions. Briefly, the PCR products were prediluted 20-fold with 10 mM Tris-HCl buffer pH 8.0. One μL of the diluted sample was separated with 39 μL Sample Loading Solution (Beckman Coulter) and 0.4 μL DNA size standard-400 (Beckman Coulter) on the GenomeLab Separation Capillary Array of the CEQ8800 (Beckman Coulter) using the Frag-3 separation program (denaturation for 120 s at 90 °C; sample injection for 30 s at 2.0 kV; separation of fragments for 40 min at 6.0 kV; capillary temperature at 50 °C). The raw data were analysed with the Fragment analysis module of the CEQ8800 software (Beckman Coulter) to estimate the size of the obtained fragments. The peak area of each fragment was exported for subsequent data processing.

***Calibration Curves.*** A calibration curve for each transcript *i* was established to calculate its relative concentration in the analysed samples from the corresponding normalized fluorescence signal values integrated over the peak areas of the respective amplified DNA fragments as separated on the capillary sequencer. To obtain the calibration curve, total RNA samples from dark- and light-treated plasmodia were mixed as described in the Results section to give an appropriate analytical standard. The mix was repeatedly diluted, two-fold in each step, to obtain a set of concentrations of total RNA ranging from 100 ng to 0.78 ng per reaction volume. Each sample was measured twice with two independent RT-PCR-reactions. The relative fluorescence signal *F_i_* (which is identical to the normalized peak area NPA) as measured for each DNA fragment *i* was obtained by normalizing its peak area relative to the peak area of the KAN^r^ control RNA which served as internal standard. The logarithm of the normalized mean values was plotted against the logarithm of the relative total RNA concentration as obtained in the dilution series [7]. By taking the logarithm for the base 10, a straight line in the form log*F_t_* = *a_i_* ⋅ log[RNA]*_tot_* + *b_i_* was computed by linear regression for the *n* data points of each fragment *i* (1^st^ linear regression).

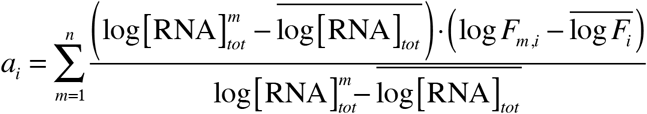

The slopes *a_i_* obtained for all gene-specific fragments in the 1st linear regression were averaged and the data points for each gene-specific fragment *i* were again fitted to a straight line by a 2nd linear regression that computes the intercept with the ordinate 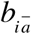 which is obtained for the average slope 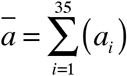 according to

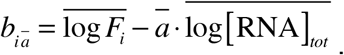

Fitting parameters for the 1^st^ and for the 2^nd^ were stored for documentation and quality control purposes. Data processing was done with the Perl scripts provided in conjunction with this paper (for details see legends to SI Figs. 4 and 5). The data points of all 35 fragments fitted the calibration curves without any systematic deviation from the straight lines. The relative concentration 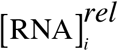 for each transcript of gene *i* was calculated on the basis of its specific calibration curve as 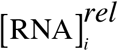 = 10^A^ with *A* = 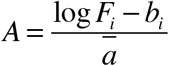

## Results and Discussion

### Kinetic model

As the concentration of the transcripts of individual genes within a given sample of total RNA may vary by orders of magnitude relative to each other, the conditions of a GeXP experiment have to be adjusted such that the amplified fragments can be reliably detected side by side when separated on the capillary sequencer. This is achieved by empirically adjusting the concentration of the gene specific reverse primers so that at intermediate physiological concentration of each transcript the amount of cDNA amplified is in the same range for all transcripts that are to be detected in one reaction tube (Fig. 2) [5–7]. Hence the concentration of the primer for the reverse transcription of certain transcripts may become rate limiting for the amount of cDNA synthesized. In the following we describe a kinetic model of the GeXP RT-PCR reaction as rational basis for evaluating the double logarithmically plotted calibration curves for the transcripts (mRNAs) to be analysed.

**Fig. 2.**
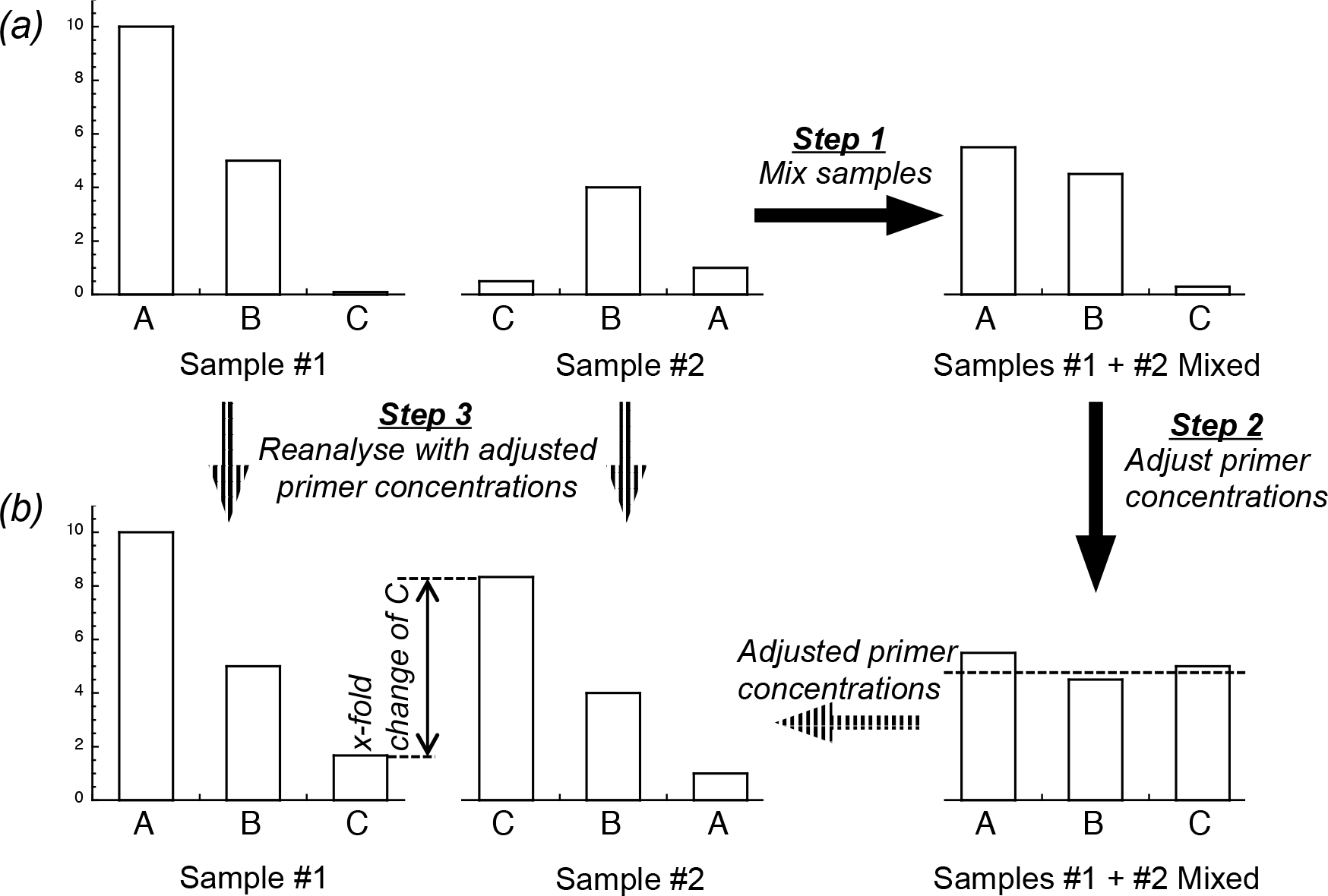
Obtaining comparable amounts of PCR products from transcripts of highly different abundances, (a) When samples of total RNA containing transcripts of genes A, B, and C that yield cDNAs of highly uneven concentrations, the amounts of some transcript-specific PCR products may be so low that differences between samples (e.g. induced *vs.* control) cannot be precisely quantified by capillary electrophoresis (as shown for transcript C). In order to allow reliable detection of the transcripts within the dynamic range of their differential regulation, the two samples (#1 and #2) are mixed (Step 1) and the concentrations of the gene-specific reverse primers (see Fig. 1) adjusted so that approximately equal amounts of cDNA fragments are obtained by RT-PCR from each of the transcripts (Step 2). Finally, the two samples, #1 and #2 are re-analysed with the adjusted primer concentrations (Step 3). Now, the x-fold change in the abundance of the transcripts of all genes can be determined due to the sufficient amount of their corresponding cDNAs (b).

Assuming that at maximum 2 % of the total RNA isolated from an eukaryotic cell is mRNA, a 10 μl RT reaction volume with the amount of total RNA as used in the experiments contains not more than 0.4 ng of mRNA. With approximately 17,000 genes expressed in a starving *P. polycephalum* plasmodium and an avarage length of a mature mRNA of approximately 1000 nucleotides [10], this is equivalent to an average concentration of 6.9 × 10^-15^ M for the mRNA of one gene corresponding to approximately 41,500 molecules per reaction volume. The smallest RT primer concentration used in our RT reaction (SI Table 1) is 2.5 × 10^-11^ M, as compared to 2.7 × 10^-8^ M for the average and 5 × 10^-8^ M for the maximal concentration, respectively. Even if an mRNA reverse transcribed with the smallest primer concentration would be 1000 times over average abundance, a concentration of 6.9 × 10^-12^ M mRNA of this gene would be reverse transcribed at 2.5 × 10^-11^ M RT-primer concentration. Hence even under these conditions the primer concentration would be in excess to the concentration of the template. Accordingly, the primer concentration for most if not all of the amplified fragments is expected to be in very large excess as compared to the template concentration and under these conditions the concentration of the reverse primer will not change significantly in the course of the RT reaction.

It has been shown that binding of a primer to its template during the reassociation (renaturation) of DNA follows the rate law of a second order reaction [11, 12]. This suggests that the rate limiting step for the reassociation is the nucleation reaction during which few bases of one strand bind to their fitting counterparts on the other strand. The subsequent zipping reaction must occur fast ascompared to the nucleation event of base pairing because otherwise a first order instead of second order kinetics would be observed. This conclusion is plausible as the local concentration of fitting bases greatly increases if two strands are brought in spatial proximity and appropriate orientation as a consequence of the zipping reaction.

The concentration of the RT-primer for the transcript of each gene of interest has to be adjusted to attenuate the relative amount of DNA amplified from its RNA during the GeXP RT-PCR reaction while the concentration of the RT-primer is in large excess as compared to the reverse transcribed mRNA (see above). We accordingly assume that primer binding

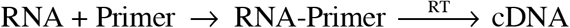

is the rate-limiting step of reverse transcription (RT). Once priming of the RT reaction has occurred, the polymerization reaction of cDNA synthesis is equally fast for all 1st strand cDNA species concurrently synthesized in the reaction mix.

Note that only a tiny fraction of the low molecular weight reactants is consumed during the RT reaction such that their concentrations do hardly change during the experiment. Based on the assumption of a rate-limiting bimolecular reaction of RT-primer binding to the RNA template, we expect an empirical rate law for the time-dependet decay *d*[RNA_*i*_]/*dt* of the concentration [RNA_*i*_] of the transcript of each gene *i* according to

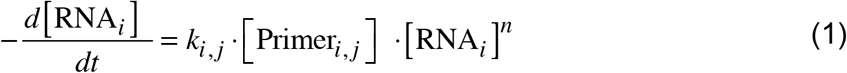

where *k_i,j_* is the primer-specific rate constant for hybridization of the primer to its complementary RNA sequence, [Primer_*i,j*_] is the concentration of primer *j* of given sequence and length, complementary to the corresponding sequence stretch of the mRNA of gene *i*, and *n* is the empirical order of the reaction. As the concentration of the primers in the RT reaction is in a sufficiently high excess over that of each gene-specific RNA (see estimation above), the primer concentrations can be approximated as remaining constant during the reaction. By defining the pseudo first order rate constant 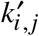 = *K_i,j_* ⋅ [Primer_*i,j*_] we obtain the empirical first order rate law

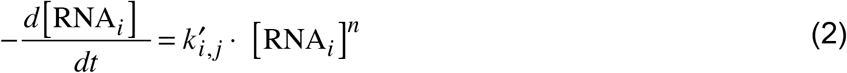

where the value of 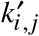 depends on the primer concentration that had been appropriately adjusted by the experimenter. Separation of the variables and integration over the reaction time *t* and the change in RNA_*i*_ concentration yields

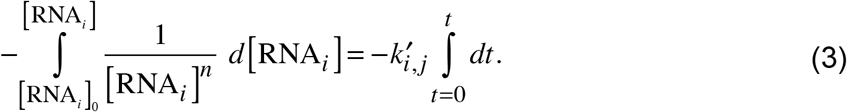

In the simplest case of a first order reaction where *n* = 1, [RNA_*i*_] decays exponentially during the reverse transcription reaction of length *t* starting from its initial concentration [RNA_*i*_]_0_ according to

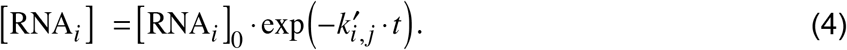

Alternatively, if the reaction order is different from one (*n* ≠ 1), the integration yields

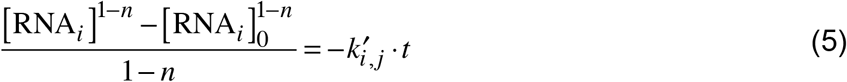

or

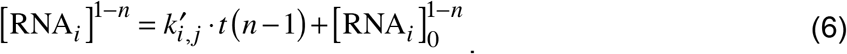

and hence [RNA_*i*_] decays according to

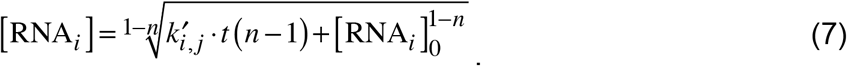

In both cases, for *n* = 1 and for *n* ≠ 1, the concentration [cDNA_*i*_] of gene-specific first strand cDNA, synthesized during the enitre RT reaction is

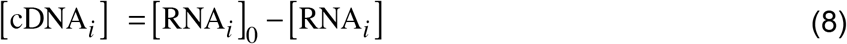

The concentration of the gene *i* -specific cDNA fragment, 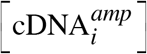 which has been amplified by PCR from the first strand cDNA 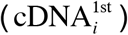

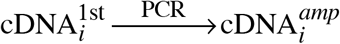

with the help of the universal primer pair (see Introduction) is expected to be

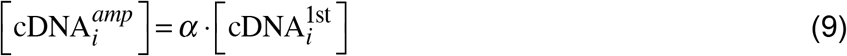

and according to eqn (8) dependent on the initial concentration of the gene *i* -specific transcript

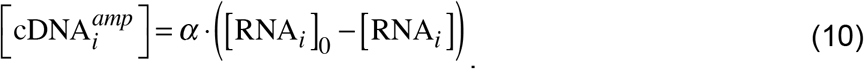

Overall amplification *α* of each 1st strand cDNA molecule after *c* cycles of PCR is expected to be *α* ≤ 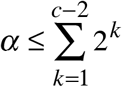 2^*k*^ because the first PCR cycle is required to synthesize the second strand on the cDNA, leaving *c* − 2 cycles for DNA amplification while a new second strand product can be synthesized from the first strand cDNAs during each cycle of PCR. Each newly synthesized second strand cDNA molecule then starts a new exponential cascade of amplification during the remaining cylces of PCR. With eqns (4) and (7), the concentration 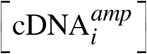 of the gene *i* -specific cDNA fragment for *n* = 1 is

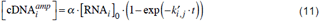

and for *n* ≠ 1

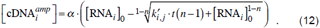

Calculating the concentration of cDNA 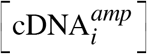 amplified from the gene *i* -specific mRNA as a function of the relative initial concentration [RNA_*i*_]_0_ in the RT reaction according to eqns (11) and (12) yields apparently linear relationships in a double logarithmic plot for empirical reaction orders of *n ≤ 1* (SI Fig. 1). The concentration of any gene-specific mRNA present in the RT-reaction is proportional to the concentration of total RNA [RNA_*tot*_]_0_ which is subjected to reverse transcription. Accordingly, a double logarithmic plot of the concentration of specific cDNAs amplified from the transcripts of different genes will yield a series of parallel shifted lines of equal slope and correspondingly different intercepts with the ordinate. The kinetic model with eqns (11) and (12) further predicts how the concentation of cDNA fragments depends on the initial concentration of the gene-specific reverse primer used for the RT reaction (SI Fig. 2).

### Experimental validation of the calibration procedure

The kinetic model of the GeXP RT-PCR predicts that for a set of transcripts one obtains a series of parallel shifted lines when the concentration of amplified gene-specific cDNA fragment is double logarithmically plotted versus the total RNA concentration used in the assay. For experimental validation of this prediction we used *Physarum polycephalum* total RNA isolated from plasmodial cells.

Starving plasmodial cells can be induced to sporulation by a brief pulse of farred light which is sensed by a phytochrome photoreceptor [10, 13, 14]. The cellular commitment to sporulation is associated to the differential expression of developmentally regulated genes [15]. Based on transcriptome data [15, 16] we have previously designed gene specific primers for the quantification of the transcripts of a set of 35 genes with a single GeXP RT-PCR reaction. The concentrations of the gene-specific reverse primers were attenuated (adjusted) so that amplification of each of the 35 cDNAs yields a sufficient amount of each fragment to be resolved side-by-side during electrophoretic separation (Fig. 2; SI Table 1) [6]. The set comprised genes that are up- or down-regulated at different strengths as well as genes with constant or almost constant expression level that can serve as a reference.

#### Fragment quantification by capillary electrophoresis

For the inital validation of the performance of the electrophoretic quantification of cDNA fragments, we used two samples of RNA, one prepared from uninduced plasmodial cells (controls) and one from far-red light induced plasmodial cells harvested at 6h after the far-red pulse. The induced cells were committed to sporulation but did not yet show any morphogenetic alterations. In the cells of the two groups, a number of genes are up- or down-regulated [6]. A known amount of spike-in RNA was added to the two samples of total RNA isolated from the cells of the two groups and the cDNAs of the 35 transcripts were amplified by GeXP RT-PCR. Subsequently, a DNA size standard was added to each of the two samples and the fluorescence tag-labelled fragments were separated electrophoretically on the CEQ 8800™ 8-capillary sequencer (see Fig. 1C). The fragments were quantified through their peak area and the amount of each gene-specific fragment was normalized relative to the amount of fragment amplified from the spike-in RNA with the help of the Perl scripts (Supplementary Information) to give the normalized peak area (NPA). In order to test the performance of the electrophoretic separation process and the fidelity of the peak quantification software, we mixed the two cDNA samples from light-induced and control cells at different ratios to obtain a series of mixtures with different concentrations of light-regulated cDNAs. Each mixture was separated on the CEQ 8800™ capillary sequencer and the normalized peak area (NPA) determined for three chosen pairs of fragments that were amplified from an up-regulated and one down-regulated transcript, respectively. For the three pairs of genes the ratio of the normalized peak areas for the corresponding two fragments was plotted against the relative concentration of the two samples in the mixture. In the three pairs of fragments the normalized peak area for each fragment was proportional to its relative amount in the mixture (SI Fig. 3), confirming that the peak detection algorithm of the software delivered with CEQ 8800™ capillary sequencer worked correctly.

#### Calibration curves for the 35 transcripts

In order to obtain calibration curves for each of the 35 transcripts, we mixed two RNA samples (from light-induced and control cells) in a 1:1 ratio in order to obtain a sample that contained even the differentially regulated transcripts at intermediate concentration (Fig. 2). This RNA mixture was serially diluted and, after addition of spike-in RNA to each sample of the dilution series, the cDNA fragments as amplified by the GeXP multiplex-RT-PCR were separated electrophoretically. The normalized peak area determined for each cDNA fragment was plotted versus the concentration of total RNA. The data points obtained for all fragments were fitted to straight lines in the double logarithmic plot. The slopes as obtained by the 35 fits were averaged and the best-fitting intercept for each of the 35 curves was calculated. Accordingly, this procedure gave a series of fitted curves that only differed in being shifted with respect to the abscissa (Fig. 3). As predicted by the kinetic model, for none of the genes any systematic deviation of the data points from the fitted straight line was seen (Fig. 3). Hence, all fragments were amplified with the same efficiency by the universal primers. The parallel shift relative to the abscissa, according to the kinetic model, is due to the different absolute concentrations of the mRNAs and to the different amounts of cDNA produced during the RT reaction at the particular concentration of the RT-primer. Noteworthy, the slopes of the curves clearly were smaller than 1 as observed in most experiments performed in our lab. A slope of 1 would be expected if the cDNA synthesis occurred with first order kinetics as assumed in eqn. (11). The kinetic model therefore suggests that the RT reaction occurs with a reaction order of *n ≤* 1 (eqn (12)). The value of *n* observed in each particular experiment might slightly vary depending on the experimental conditions.

**Fig. 3.**
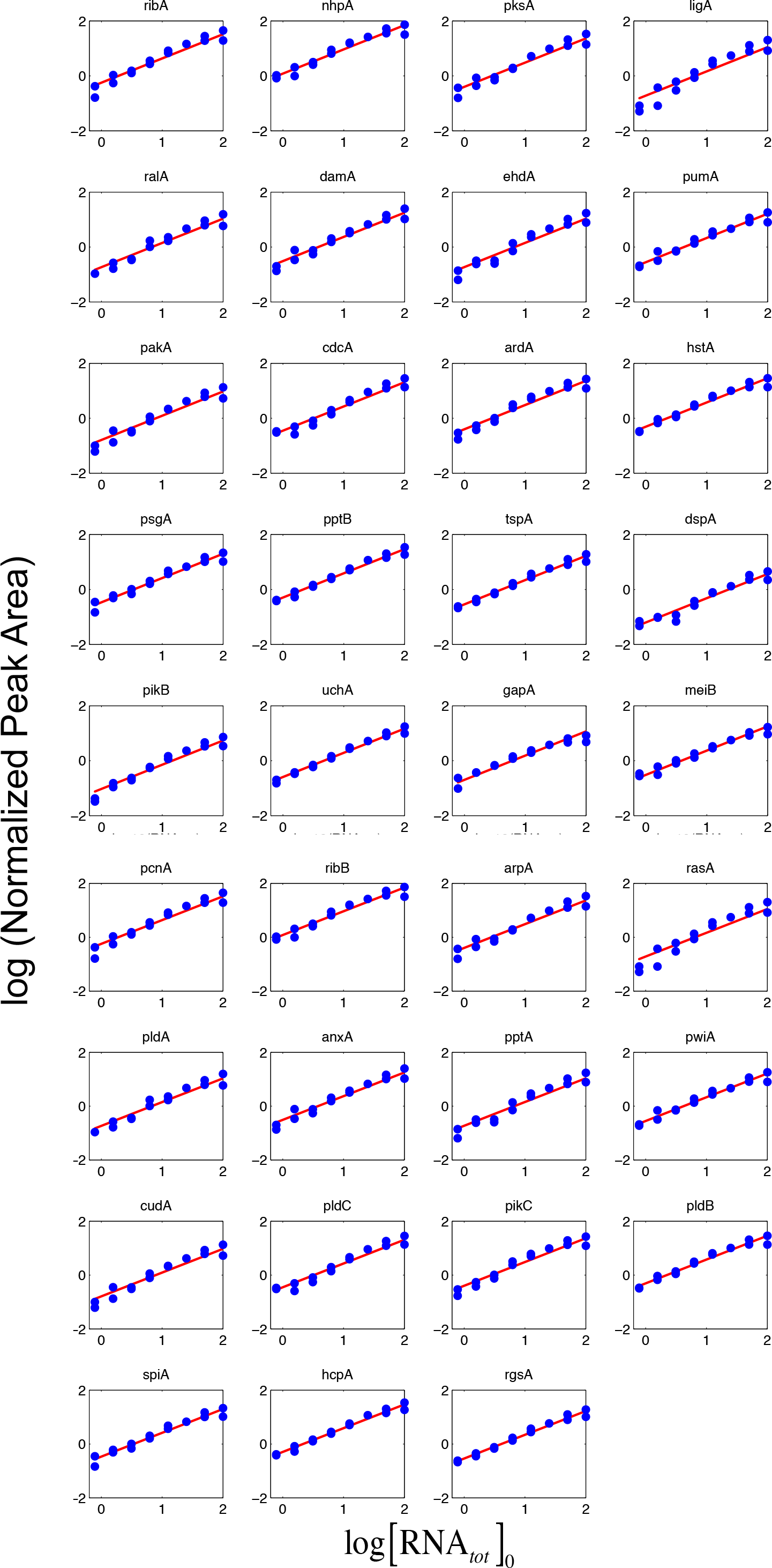
Calibration curves for the transcripts of 35 genes as amplified by GeXP RT-PCR in the same tube. A sample of a total RNA pool from light-induced and control plasmodial cells (see Fig. 2) was serially diluted and the 35 cDNA fragments quantified after capillary electrophoretic separation. Each RNA concentration was analysed twice and the two resulting data points, normalized peak area (NPA) versus concentration of total RNA, displayed in a double logarithmic plot (logarithm to the base 10). All data points were fitted to a straight line of the same slope as described in Materials and Methods. For names of genes see SI Table 1.

#### Efficiency of concurrent amplification of different transcripts in one sample

Since cDNA fragments must be of different lengths in order to get separated during capillary electrophoresis and the efficiency of amplification might depend on the fragment length, we wished to confirm that the amount of cDNA amplified for any given transcript is independent of other fragments that are concurrently amplified in the same sample. Therefore we mixed the RNA sample from the control cells with the sample from the light-induced cells taking different relative amounts and estimated for each gene how the normalized peak area changed with the relative amount of the RNA from light-induced cells or from the dark controls (Fig. 4). For each fragment, the normalized peak area and hence the amount of amplified cDNA was a linear function of the mixing ratio with a slope depending on whether genes were up-regulated or down-regulated. Curves for genes that are not regulated in response to the photo-stimulus fitted to a straight line that was parallel to the abscissa (Fig. 4). Hence, the amount of amplified DNA for each gene was found to be proportional to the concentration of the corresponding mRNA in the sample. Likewise, amplification of the cDNAs of those genes that are not or rarely differentially regulated (*dspA, spiA, tspA, etc*.) confirms that the amplification efficiency indeed does not depend on the concentration of other (differentially regulated) mRNAs that are concurrently amplified in the sample.

**Fig. 4.**
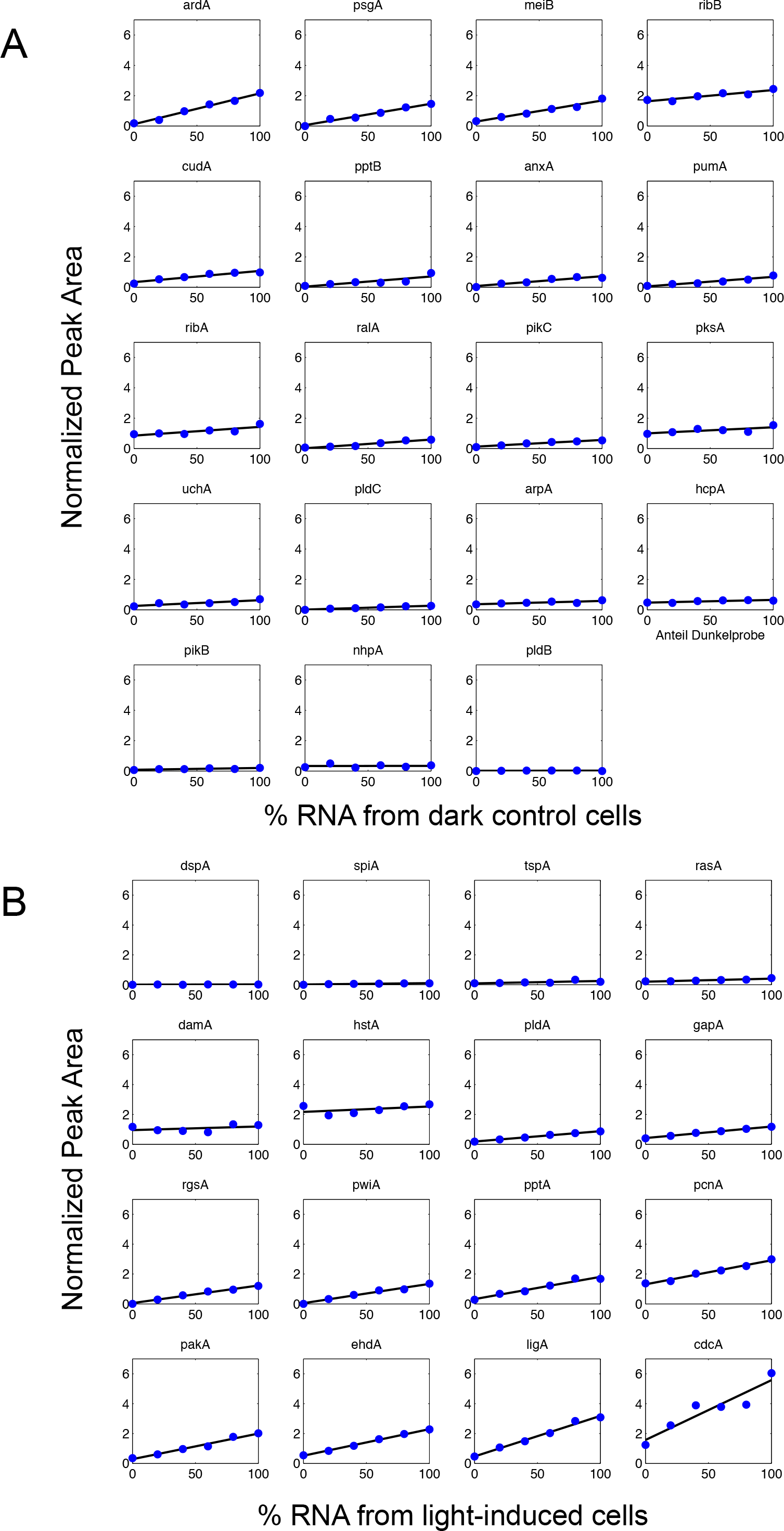
Efficiency of concurrent amplification of different transcripts in one sample. Two RNA samples isolated from light-induced and from dark control cells, respectively, were mixed at different ratios. Transcripts were quantified and the normalized peak area (NPA) of their specific fragments was plotted against the relative concentration of the RNA sample from the dark control cells (A) or from the light-induced cells (B). Data points of down-regulated (A), up-regulated (B), and not regulated genes fitted to straight lines, indicating that the amount of cDNA amplified by RT-PCR is proportional to the concentration of each individual transcript in the sample and independent of those transcripts the cDNAs of which are co-amplified.

An experimental result consistent with the computational predictions of the kinetic model of the GeXP RT-PCR (eqn (12)) was obtained by changing the concentration of one primer while the concentration of all other primers was kept constant. Until saturation, the normalized peak area increased with the RT-primer concentration, indicating that the reverse primer was rate-limiting even though its concentration was high relative to the concentrations of the other RT-primers in the mix (SI Fig. 2).

Taken together, these experimental results confirm that all fragments are amplified with the same efficiency during PCR with the universal primer set regardless of their sequence or the length of the amplification products. Amplification of individual fragments is not influenced by other fragments that are concurrently amplified in the sample. This confirms (at least for the primer set chosen and for the experimental conditions applied) that the relative changes in the mRNA concentration of any fragment robustly translate into relative changes in the amount of detected amplified cDNA during a GeXP multiplex-RT-PCR experiment. Certainly, equal amplification efficiency for all cDNAs quantified within one essay should be verified for any new set of primers and any new set of transcripts to be analysed. The Perl scripts provided as supplementary material (see below) facilitate this task.

### Quantification of differentiation marker genes in plasmodial cells before and after induction of sporulation

By quantifying the light-induced expression of differentiation marker genes in *Physarum polycephalum* plasmodial cells, we present a workflow including the application of Perl scripts for calibration and data evaluation.

#### Stimulation of cells, sample preparation, and biological replicates

Plasmodial cells were starved for six to seven days to gain sporulation competence. Sporulation-competent plasmodial cells, each spreading the surface of a Petri dish were stimulated with a short pulse of far-red light for the induction of sporulation. Before and at different time points after application of the stimulus, samples of each plasmodial cell were harvested and frozen in liquid nitrogen for the subsequent isolation of RNA. The remainder of the plasmodium was further incubated overnight to verify whether the developmental decision to sporulation had occurred [17]. This approach gave a collection of time series where each time series consisted of samples taken from the same individual plasmodial cell. In a control experiment, light stimulation was omitted while samples of individual plasmodial cells were taken correspondingly. From each frozen sample, total RNA was isolated and the expression pattern of the set of 35 genes (SI Table 1) were quantified as described above.

#### Data processing workflow and Perl scripts

The electrophoretic separation of the GeXP multiplex RT-PCR samples was performed in the CEQ 8800™ Genome Analysis System run via the so-called Control Center, a software package for operating the system and for storing the data. All further data processing steps were performed by Perl scripts as described in the Supplementary Information (see also SI Figs. 4 and 5).

As a standard quality control for the electrophoretic separation, the electropherogram and the detected fragments were graphically displayed and visually inspected calling the respective scipts (SI Fig. 4). All further data processing steps were executed using the .ceqfrag files that list peak height, peak area and length (in nucleotides) of each fragment. The Perl scripts *gep_makestd* and *gep_calcamount* perform a batch evaluation of these files to generate the standard curve and to calculate the relative transcript abundances from the measured data, respectively. This is achieved by taking the fitted mean slope and the corresponding intercept of the standard curve for each transcript into account. The final output is a table in the form of a file with tab-separated values that contains for each quantified fragment the fragment ID number, the length (nt’s), peak area, normalized peak area (NPA), the relative amount of cDNA, and the detection limit of each fragment during the respective separation run. Two values, relative amount of cDNA and detection limit, are calculated based on the calibration curves. We routinely made a new standard curve for each kit in order to account for possible variations in the performance of the kit components. Details on data processing and on the determination of the transcript-specific detection limits are given in the legend to SI Figs. 4 and 5. Samples of all file types used are available together with the Perl scripts (for details see Supplementary Information). With the help of the two Perl scripts, the data processing for the generation of the calibration curves and for calculating the amounts of cDNA for each fragment and sample becomes fully automatic due to the batch processing of the files. We finally store all values for calibration curves as well as for all other measurements in an SQL database for documentation and further data processing and evaluation.

#### Light-induced gene expression and technical replicates

For visual comparison of the light-induced differential regulation of the differentiation marker genes in individual plasmodial cells, the expression values of the 35 genes were comparatively displayed for each sample (Fig. 5; for the complete data set see SI Fig. 6). Each data point of a time series originates from material taken from the same plasmodial cell, allowing to compare time series between individual cells. Strongly up- or down-regulated genes are clearly evident while the expression of *ribB* was almost constant. Accordingly, *ribB* might be used as a reference gene to normalize the expression values in each sample in order to correct for pipetting errors or inaccuracies in the quantification of the total RNA samples analysed. Nevertheless, we displayed all expression values as they were obtained, i.e. without normalization. However, the five expression values for each gene in a time series taken from a given cell were then normalized to the maximal value to reveal the percental changes as a function of time in order to highlight the relative changes that occurred as a function of time.

**Fig. 5.**
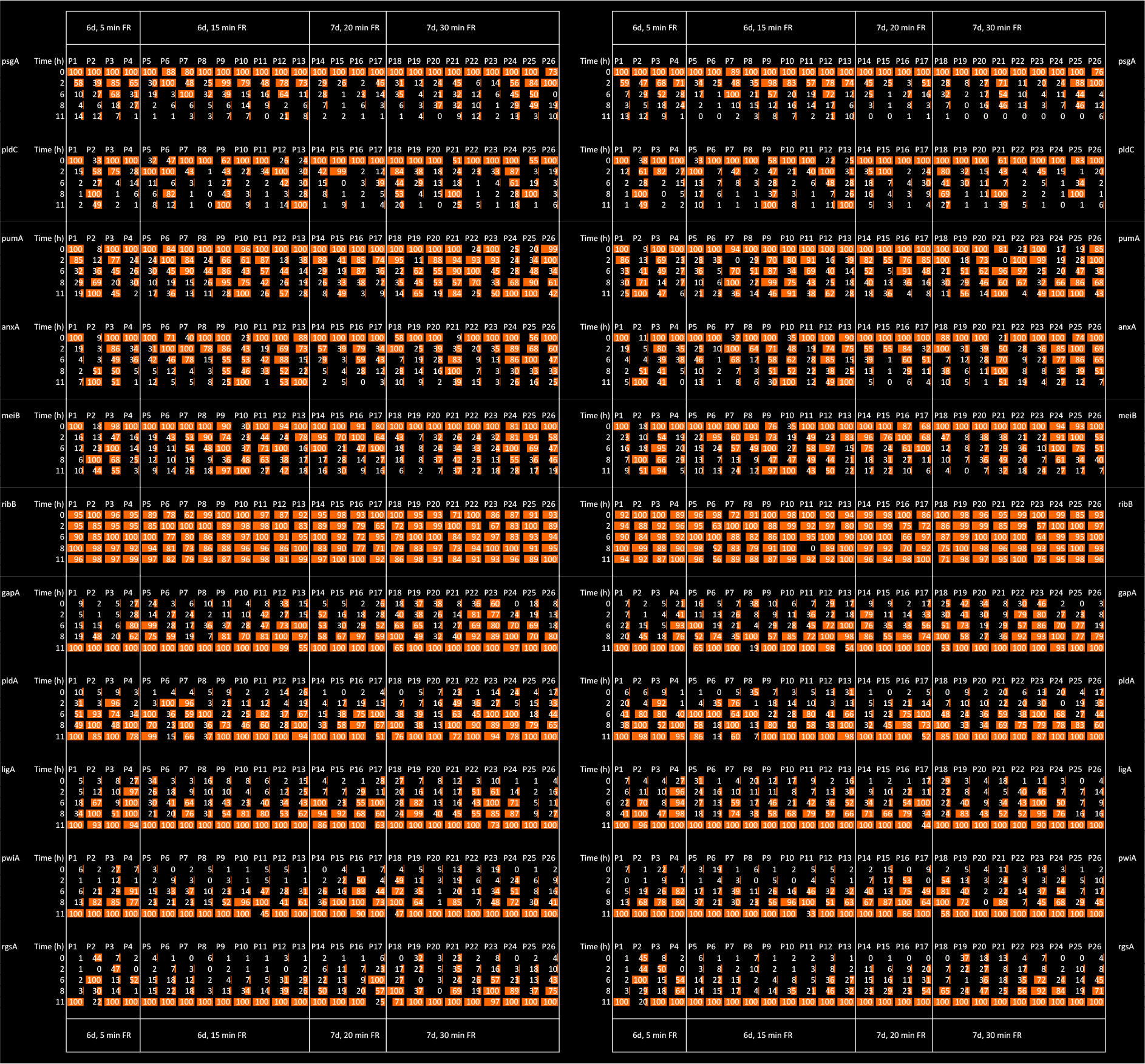
Comparative display of single cell gene expression time series in response to a sporulation-inducing far-red light stimulus. Individual plasmodial cells (P1 to P26) starved for 6 or 7 days to gain sporulation competence were exposed to a far-red light pulse for 5 to 35 minutes. Before and at several time points after the light pulse, samples were taken from each plasmodium and the gene expression pattern analysed to give a time series for each individual cell. The relative changes of expression values were normalized to the maximal value within the time series and displayed side by side for each plasmodium and gene. The gene expression pattern in each of the RNA samples was analysed twice with independent RT-PCR reactions and based on separate calibration curves. The first measurement and the technical replicate are displayed in the left and in the right panels, respectively.

In order to estimate how reproducible the measurements are, the RNA samples were measured again in an independent GeXP RT-PCR experiment which was evaluated based on a fresh calibration curve. The results of the technical replicate were displayed side by side with the first measurement (Fig. 5; SI Fig. 6) and revealed that the time series gave similar results. For a quantitative evaluation, the reproducibility of the measurements was analysed with the help of a frequency distribution. For each single expression value obtained in the first and in the second measurement (the technical replicate) for any given sample and gene, the ratio between the two values was calculated and the absolute frequency of the log2 (ratio) displayed in the form of a histogram. For the experiment performed with light-induced plasmodia (Fig. 6A) the variation was considerably higher than for a second experiment perfromed with plasmodia that had not been exposed to any light (dark controls; Fig. 6B). For the light-induced plasmodial cells, 77 % of the values varied by a factor of two or less and 9 % of the values by a factor of four or more. One reason for the high variation observed in the light-induced plasmodia might be that genes are strongly up- or down-regulated. Two measurements of a transcript, the concentration of which is close to the detection limit, may be inaccurate and therefore give a relatively high value for the ratio between first and second measurement. In contrast, the values measured in the dark controls were highly reproducible (Fig. 6B).

**Fig. 6.**
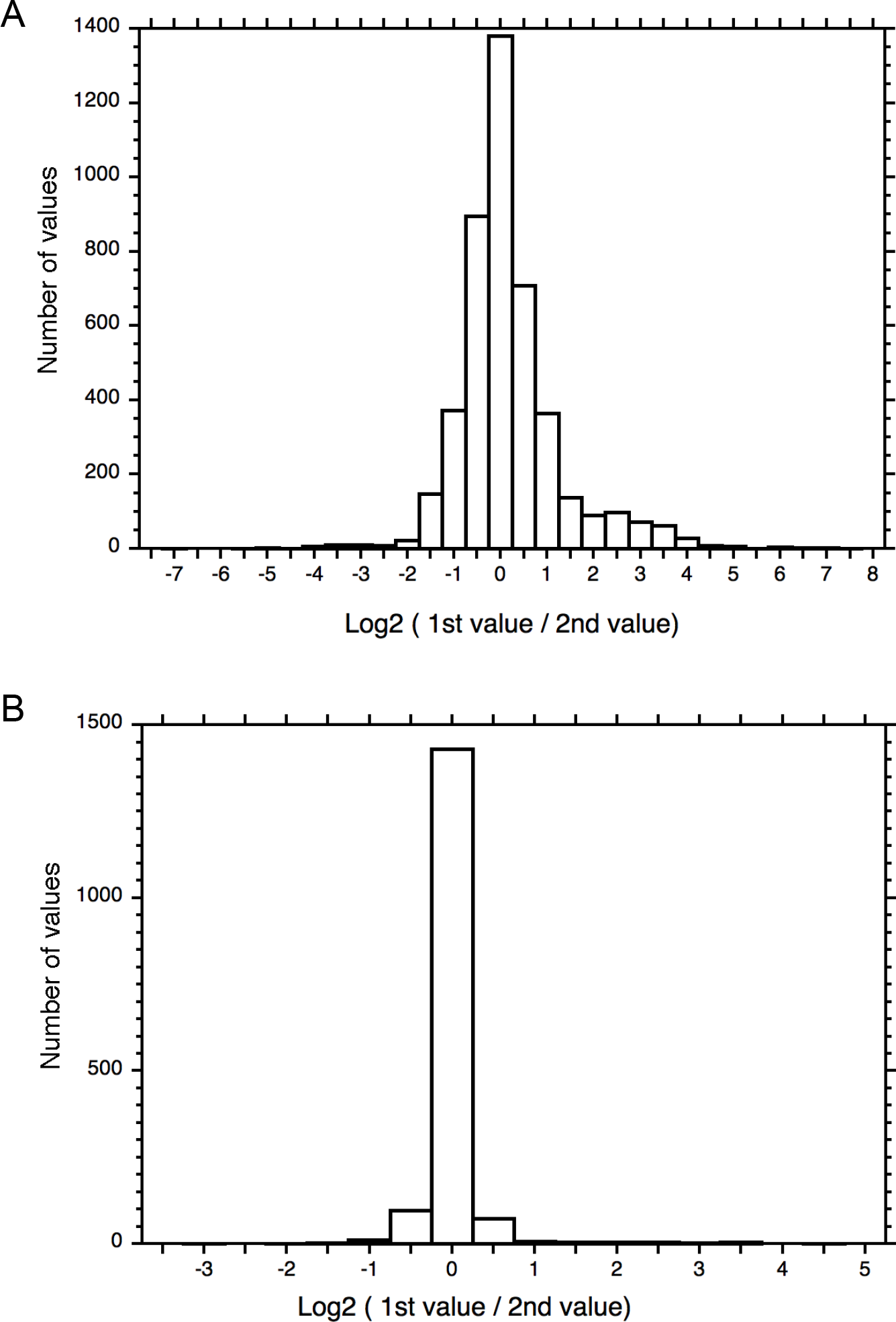
Frequency distributions of X-fold discrepancies between the gene expression values obtained by two repeated measurements. For each RNA sample and for each of the 35 genes analysed the Log2 of the ratio of the two independently measured expression values was calculated and the number of occurrences for each interval displayed in the histograms. The distributions were shifted on the abscissa so that the most abundant class was centered at an Log2 value of zero. Expression values that were below the detection limit were not considered. Panel (A) shows the distribution of 4447 values as displayed in Fig. 5 and panel (B) shows the result of the technical replicate of a set of time series (1643 values) measured from plasmodia that had not been exposed to any light stimulus (dark controls).

## Authors' contributions

All authors contributed to the development of the method. MH and WM conceived of the study and participated in its design and coordination. BW, VR and PM performed measurements. MH designed the data processing workflow and wrote the Perl scripts. WM developed the kinetic model. PM revised the model and performed model calculations. PM and BW prepared some of the figures. WM wrote the manuscript. All authors read and approved the final manuscript.

## Acknowledgments

We thank PD Dr. Marcus Hauser for discussion and valuable suggestions, Dr. Xenia Hoffmann for help with the literature search and Drs. Manfred Souquet and Marcus Hauser for critical reading of the manuscript. This work was partly funded by the Deutsche Forschungsgemeinschaft (DFG; MA 1516/10-1). BW and PM were financially supported by the International Max Planck Research School through the Research Center Dynamic Systems, Magdeburg, funded by the state of Saxony Anhalt.

## Supplementary Information

Supplementary Information is available through the online version of this article.

## Quantifying 35 transcripts in a single tube: Model-based calibration of the GeXP RT-PCR assay

Pauline Marquardt, Britta Werthmann, Viktoria Rätzel, Markus Haas and Wolfgang Marwan

## Supplementary Information

**SI Fig. 1.**
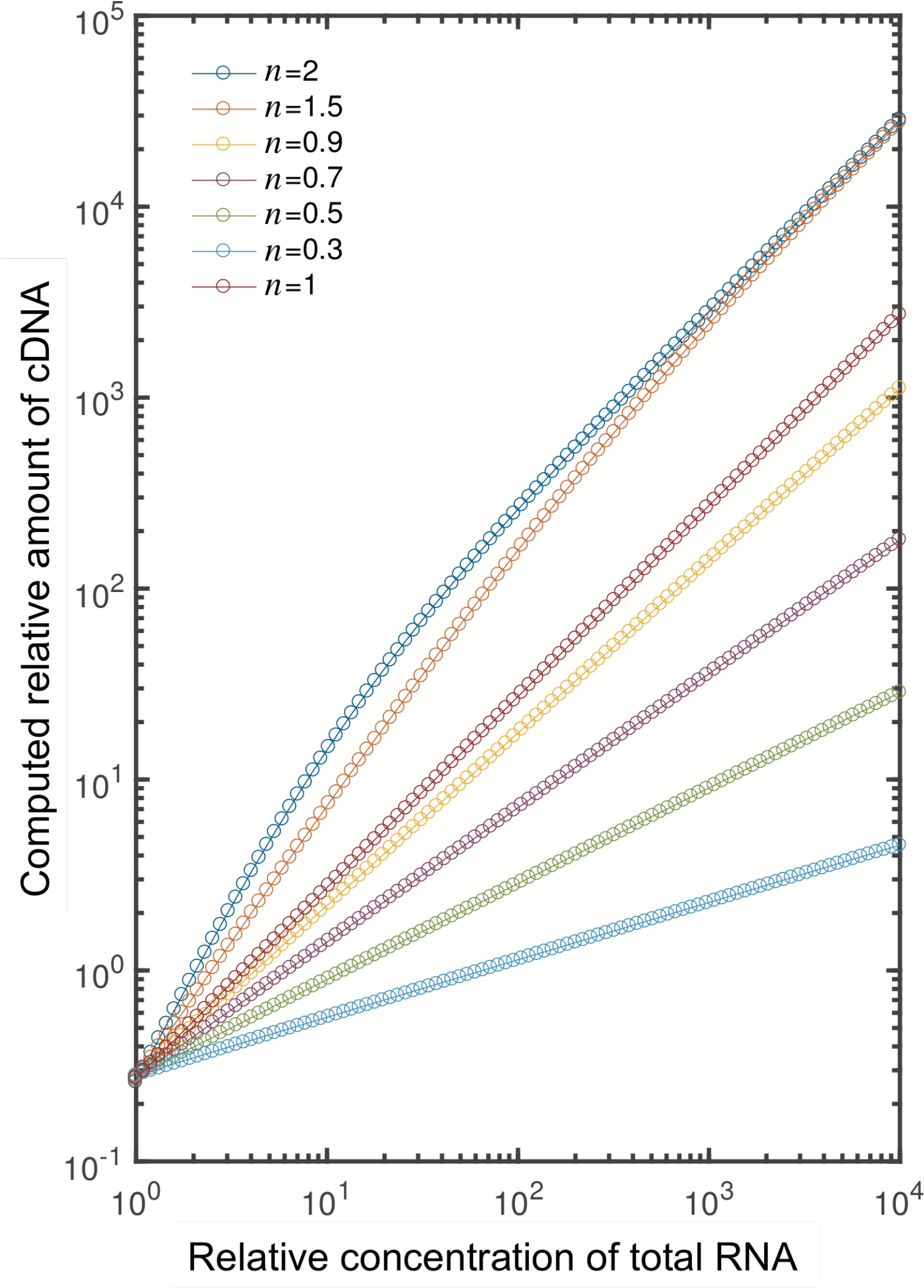
Calibration curves as predicted by the kinetic model of the GeXP RT-PCR reaction for different empirical reaction orders *n* as indicated in the inset. The relative concentration of cDNA synthesized during the RT reaction was computed as a function of total RNA concentration according to eqns. (11) and (12). The model predicts approxiamtely linear calibration curves for reaction orders of *n* ≤ 1 in the double logarithmic plot.

**SI Fig. 2.**
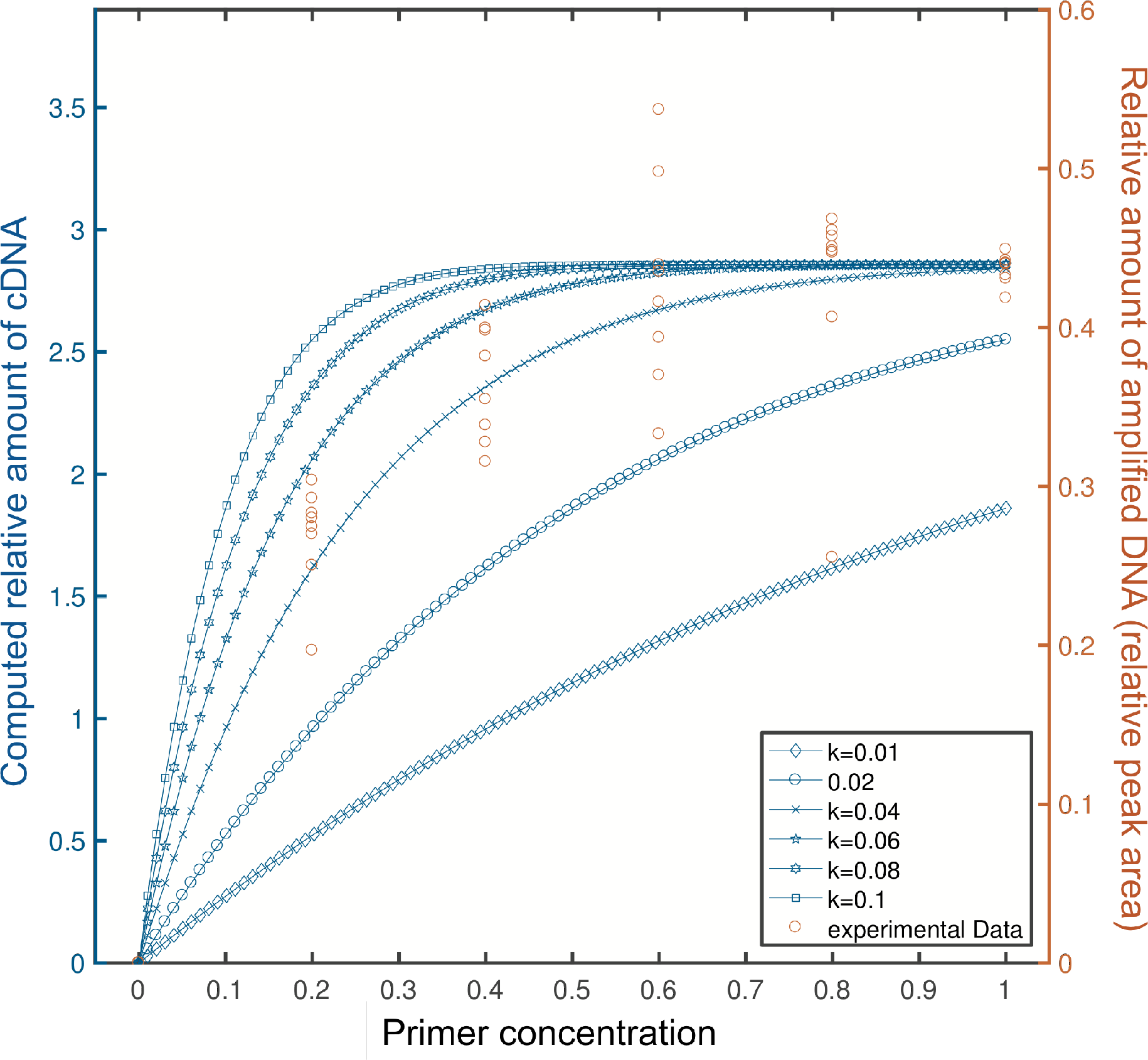
Concentration of an amplified cDNA fragment as a function of the concentration of the *pcnA*-gene-specific reverse primer used for the RT reaction. Experimental values are represented by brown symbols. Blue symbols represent computational results for a reaction order of *n* = 0.9 as obtained from eqn. (12) for different values of the primer-specific rate constant *k'*. The relative primer concentration of 1.0 corresponding to 50 nM was used in the RT reaction of the standard GeXP experiments.

**SI Fig. 3.**
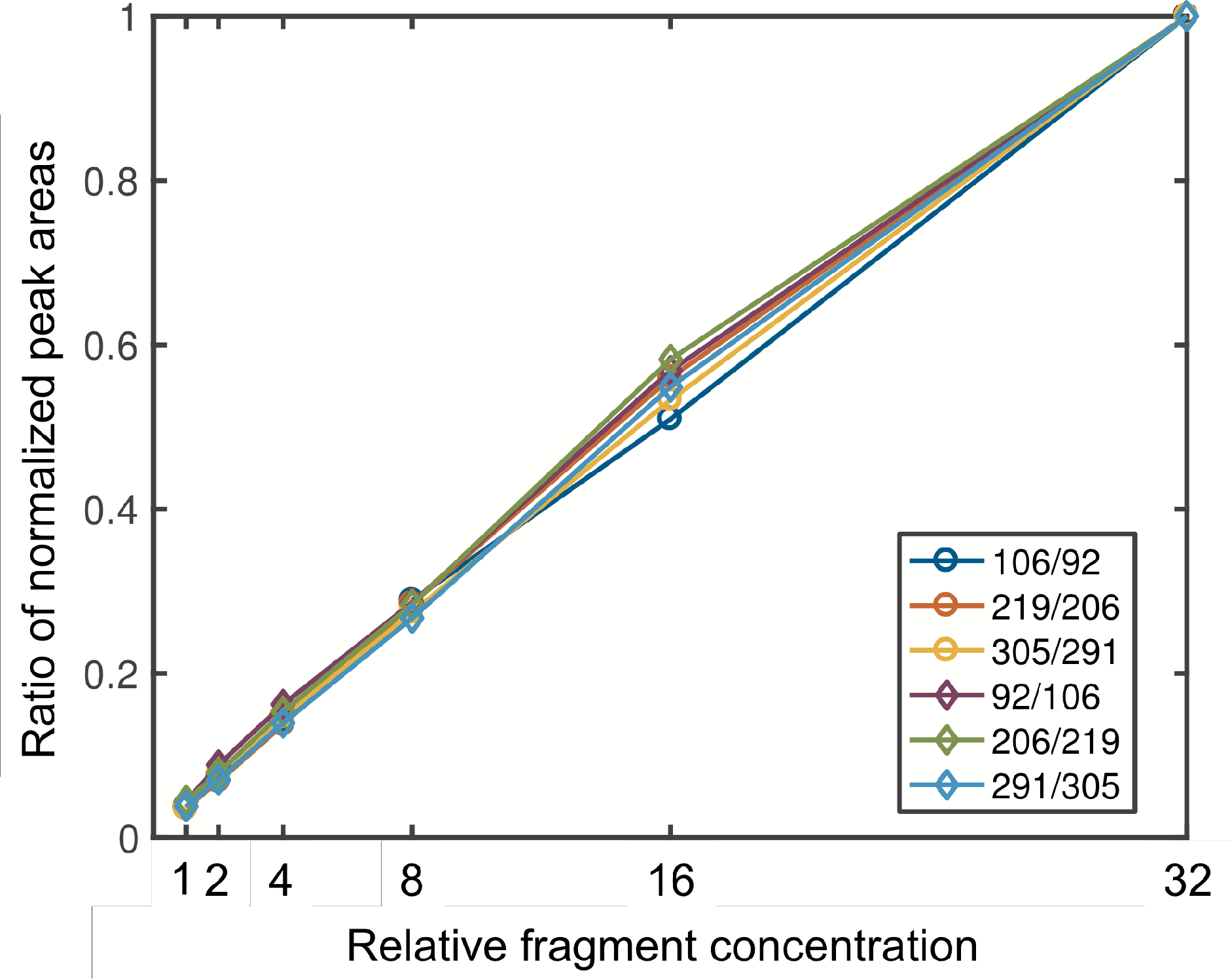
Calibration of fluorescent DNA fragment quantification by the CEQ 8800 Genetic Analysis System. Two cocktails, each containing three fluorescently labelled PCR products of different lengths (as amplified from phage *λ* DNA) were prepared (cocktail #1: 92 nt, 206 nt, 291 nt; cocktail #2: 106 nt, 219 nt, 305 nt). The cocktails were mixed at different relative fragment concentrations, separated on the CEQ 8800 Genetic Analysis System, and the normalized peak area (NPA) for each fragment determined using the custom software delivered with the system. The ratio of the normalized peak areas (NPAs) measured for pairs of fragments (as indicated in the inset) were plotted against the relative concentration of cocktail #1 or cocktail #2, respectively. Values fitted to a straight line with a slope of 1, confirming that the peak area as determined by the software was proportional to the amount of DNA fragment subjected to capillary electrophoresis.

**SI Fig. 4.**
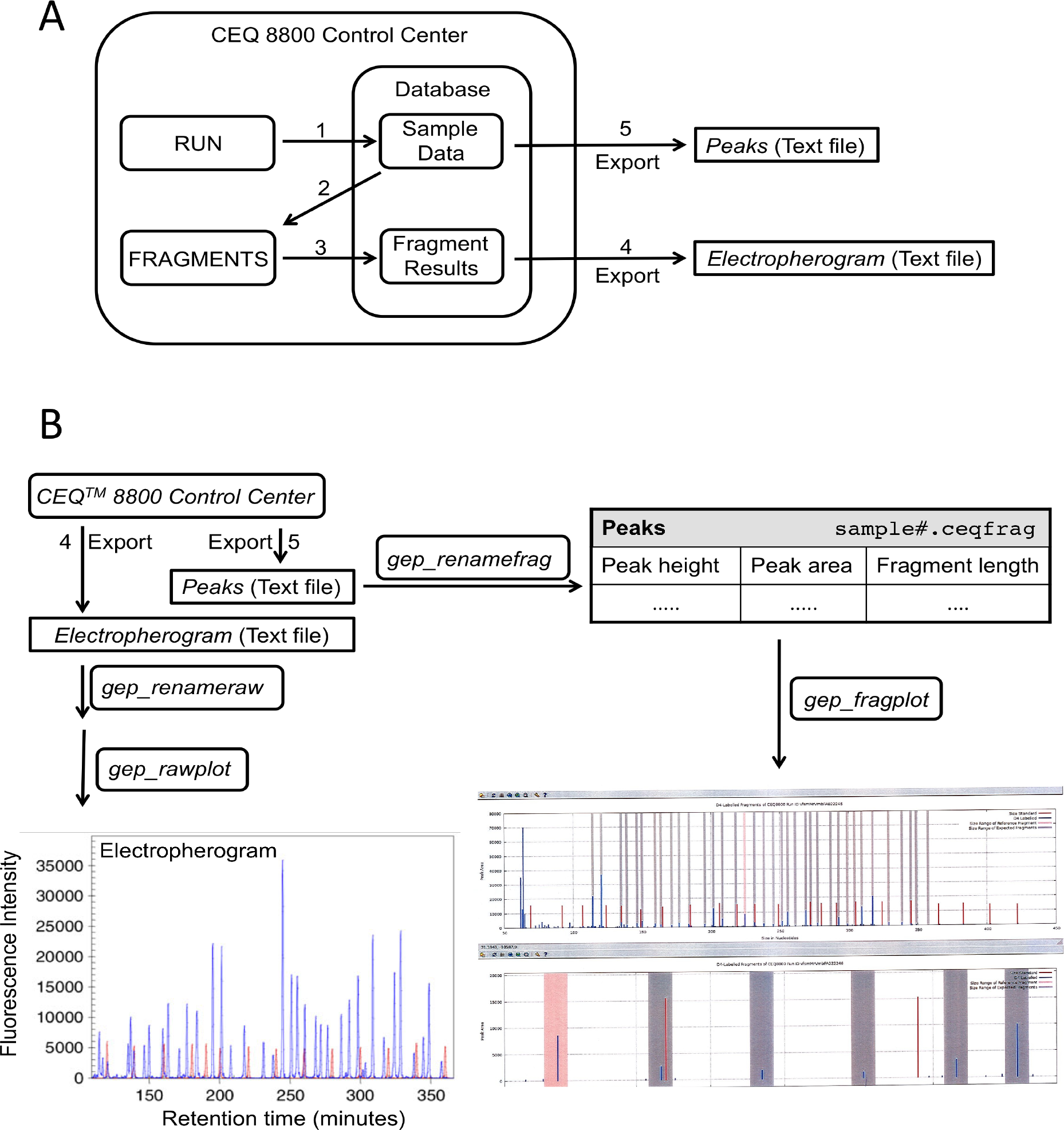
Electrophoresis and signal processing. (A) A DNA ladder of fragments of known size is added to each sample of amplified cDNAs and fragments are separated electrophoretically on a CEQ 8800 analyser. The CEQ™ 8800 software (Control Center) saves the electropherogram as Sample Data in the database (step 1). After the run is completed, the operator calls the *FRAGMENTS* module of the Control Center to generate the Fragment Results, a table which assigns the peak height, the peak area, and the calculated fragment length to each peak detected above threshold (step 2) which are also stored in the database of the Control Center (step 3). All peaks for which the peak height is smaller than 0.5% of the height of the second largest peak are considered to be below threshold. The cutoff value of 0.5% as used in this study is to be specified by the user of the CEQ™ 8800 Control Center, is kept constant for all experiments, and defines the detection limit for each peak. The *Peaks* (Fragment Results, consisting of peak height, peak area, and fragment length) and the *Electropherogram* (Sample Data, fluorescence intensity versus retention time) of each sample are exported from the database of the CEQ8800 Control Center in the form of tables as tab-separated text files (steps 4 and 5, respectively). The *Peaks* file is lateron used for calibration or fragment quantification (see SI Fig. 5). (B) For convenient analysis, the *Electropherogram* and *Peaks* files may be transferred to a separate personal computer for further data processing. The *Electropherogram* file extension is renamed to change the file extension to .ceqraw and displayed in Gnuplot by running the scripts *gep_renameraw* and *gep_rawplot*, respectively. The graphical representation of each electropherogram (panel lower left; peaks of cDNA fragments in blue, peaks of the DNA size standard in red) is visually inspected to assess the quality of the separation run before the data are further numerically analyzed. The *Peaks* file extension is renamed by the script *gep_renamefrag* as well in order to specify the file type. The panel lower right is generated by the script *gep_fragplot.* It graphically displays the detected peaks in relation to the size ranges of the expected fragments (blue) and the size ranges of the reference fragments (of the DNA size standard; red) which both are used to automatically assign the detected peaks to the respective fragments. With the help of this plot one can assess the quality of each separation.

**SI Fig. 5.**
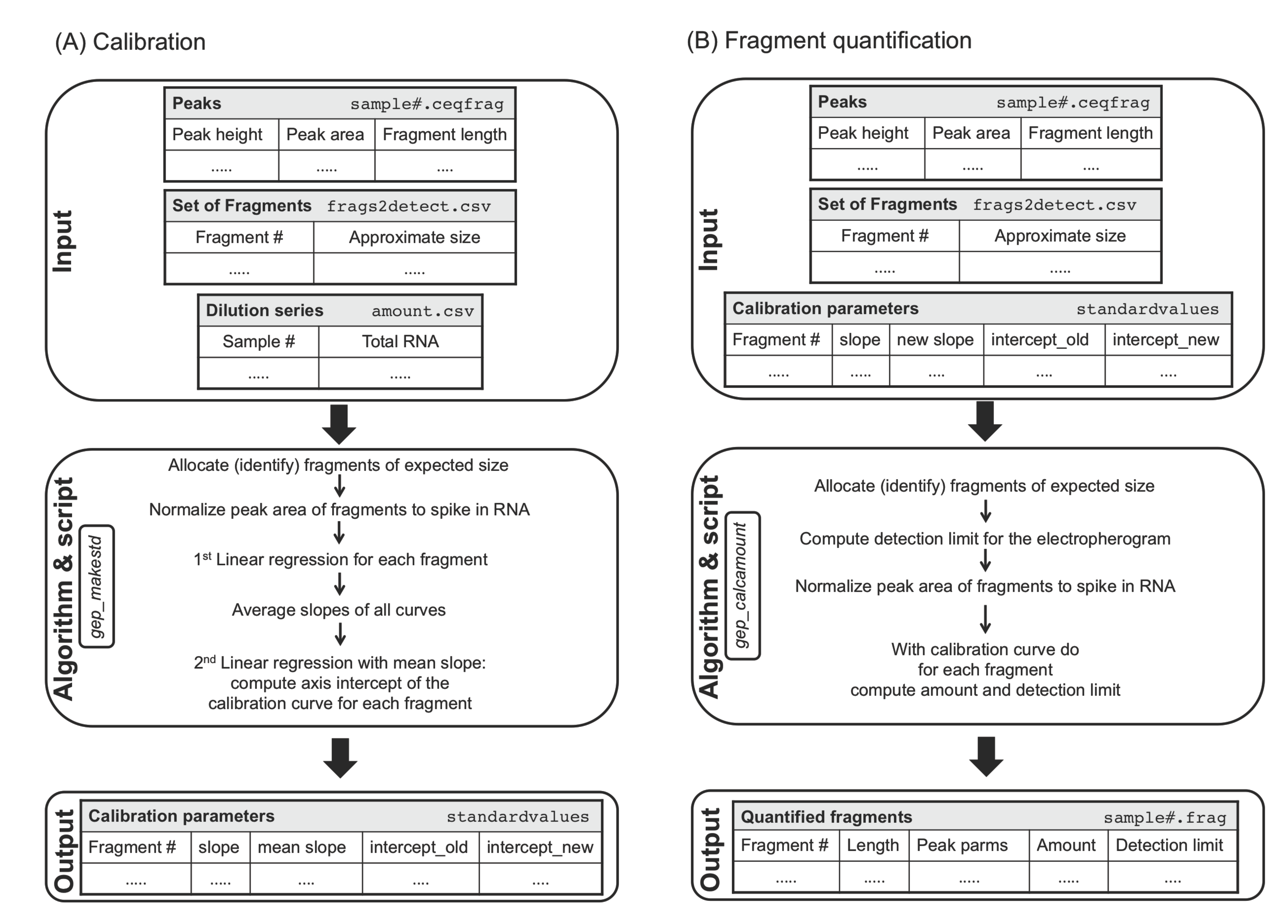
Script-based evaluation of Peaks. (A) Calibration. A calibration curve is recorded for each of the fragments to be analysed. To generate the calibration curves, an RNA sample that contains the RNAs of interest at intermediate concentration (see legend to Fig. 2 for details) is serially diluted and the diluted samples with their total RNA concentrations are entered to give the table *Dilution series* that together with the list of expected fragments (*Fragments)* serve as an input read by the script *gep_makestd.* For all fragments to be quantified, this script identifies fragment-specific peaks by assigning the largest peak from the *list of fragments* file that has a measured fragment length of ±0.8 nt of the expected approximate size as listed in the *Fragments* file for the respective fragment. The script then normalizes the peak area of each fragment to the peak area measured for the spike-in RNA. With these values the script computes for each fragment a calibration curve by performing a linear regression of the log (normalized peak area) versus log (total RNA concentration) values (1st linear regression). The slopes obtained for all expected gene-specific fragments in the 1st linear regression are averaged and the data points for each gene-specific fragment are fitted to a straight line by a 2nd linear regression that computes the intercept obtained for the average slope. The resulting curve fit parameters of each gene-specific fragment to be analysed, are stored in the *Calibration curves* file. The entries in the *Calibration curves* file are inspected line-wise for consistency of slope and intercept values obtained by the 1st and the 2nd linear regression. (B) Fragment quantification. The curve fit parameters (mean slope and intercept obtained by the 2nd linear regression) of the calibration curves (as contained in the file *Calibration curves)* are used to quantify gene-specific fragments by evaluating *Peaks* files obtained by analysing experimental samples. For each gene-specific fragment in each sample, the script *gep_calcamount* calculates its relative amount and the corresponding detection limit. The detection limit may be different for each gene-specific fragment and differ between electrophoretic separations. It is estimated taking the 0.5% value of the height of the second largest peak in the electrophoretic separation of the analysed sample and the parameters (mean slope and intercept) of the 2nd linear regression of the valid calibration curve obtained for each respective fragment. For download of the relevant scripts and example data files see Supplementary Information.

## Legends to Additional Supplementary Files

**SI Fig. 6.** Comparative display of single cell gene expression time series in response to a sporulation-inducing far-red light stimulus, first measurement and technical replicate. The figure which is provided in the form of a separate supplementary file to the online version of this article (SI_Figure 6.pdf) shows a full-length version of Fig. 5 with the values for the transcripts of all 35 genes displayed. For further details see legend to Fig. 5. Because of its size, the figure is best inspected with the help of a pdf viewer with appropriately adjusted zoom factor.

**SI Table 1.** Sequences and concentrations of primers used for GeXP-RT-PCR. The Table is provided in the form of an Excel sheet and is provided as part of the Supplementary Material to the online version of this paper.

## Protocol for the installation of Perl scripts for the GeXP data analysis workflow

### Installation under Linux

To run the software on a Linux computer, retrieve the files from https://github.com/markushaas/genexpro4ceq8800 and install them by following the install instructions that are provided together with the files.

### Installation under Mac OS X or Windows^R^

For running the Perl scripts under Mac OS X or Windows^R^ operating systems (os), we have set up a virtual Linux machine (vm), a package, in which the Linux operating system and all files necessary for the GeXP data analysis workflow (Perl scripts, example files, and ReadMe) are put in place. Download from github is therefore not necessary. Installation of the virtual machine can be easily performed by going step by step through the following protocol:

- Install "VirtualBox" from Oracle (https://www.virtualbox.org/).
- Download the vm image file we have prepared by clicking on the following link http://www.regulationsbiologie.ovgu.de/Downloads.html or by copying the link into the address bar of your browser window.
- Open "VirtualBox" and select "Import Appliance…" from the "File" menu, choose the vm image file "gep_vm.ova" you have downloaded and follow the instructions of the VirtualBox" dialogue.
- To share a folder with your host os (for shutteling files between your os and the virtual machine), create a folder with the name ".virtualboxsharedfolder". If you are using a Windows^®^ pc you may need to create the folder within a terminal window of your os as the file manager does not accept names beginning with a "." so execute cmd.exe and enter "mkdir .virtualboxsharedfolder".
- Right click on the "gep_vm" item that shows up in the VirtualBox window and select "Settings" and therein "Shared Folders". Click on the "Add new shared folder" icon, select the folder ".virtualboxsharedfolder", and enable "Auto-mount". Files in this folder can be accessed by both, your host and the operating system run by the virtual machine.
- Start the vm. Depending on the BIOS installed on your computer, you may have to enable "Intel^®^ Virtualization Technology" in the BIOS settings in order to allow you to use a vm on your pc. Refer to the manual of your hardware if necessary.
- After booting the vm open the "Introduction" file (by double clicking the icon) on the Desktop of the vm to get some help with the GeXP data analysis workflow. To adapt the resolution of the virtual screen, adjust the settings in the "view" menu of the virtual machine window.
- In case of problems with the shared folder or the screen resolution that might occur under the particular os you are working with, refer to the internet to get help.

## Footnotes

1 The Beckman Coulter Capillary Electrophoresis product line including GeXP chemistry is now distributed and supported by AB SCIEX, Landwehrstr. 54, 64293 Darmstadt, Germany. (www.SCIEX.com).

